# Effect of the shrimp farming wastes as co-feed on growth performance and digestibility of juvenile grey mullet, *Mugil cephalus*

**DOI:** 10.1101/2021.04.29.441540

**Authors:** Jorge Madrid, Zohar Ibarra-Zatarain, Jorge E. Viera-Pérez, Abelardo Campos-Ezpinoza, Emilio Peña-Messina

## Abstract

A feeding trial was carried out to evaluate the utilization of residual nutrients from a shrimp farming wastes as co-feed in different proportions in juvenile grey mullets. Four treatments were designed offering shrimp farming wastes at 0, 33, 66, and 100%. The 4 % of feed respecting the total biomass of each experimental tank was offered daily. The total feed ratio was completed with commercial feed for marine fish in the treatments where it was needed. At the end of the feeding trial, final weight, weight gain, relative weight gain, and thermal growth coefficient were reduced in the fish when increasing the amount of SFW as feed. However, the fish showed a digestive capacity to use residual nutrients up to 66 %, increasing their initial weight by up to 25 %. The increase of shrimp farming waste as feed negatively affected the whole-body proximal composition. The digestibility results showed that the fish could digest up to 41 % of the shrimp farming waste protein. Results suggest that juvenile grey mullets are capable of utilizing residual nutrients from shrimp farming waste. However, it is necessary to use an alternative feed source to induce an optimal growth performance for the juvenile grey mullets. It is also recommended that mullets be fed with formulated feed to meet their nutritional requirements to maintain the protein and lipid content of the whole-body under culture conditions.

## 1. Introduction

World aquaculture production from 2000 to 2018 sustained annual growth of 5.3 %. Total production was reported at 82.1 million metric tonnes of aquatic animals (FAO. 2020). Nevertheless, problems associated with the production increment have also arisen. The excessive supply of nutrients from the crop waste (i.e., ammonia and phosphorous), mainly formed by uneaten feed and feces, has caused eutrophication problems in the environment (Thomsen et al., 2020; Ottinger et al., 2016; Gowen, 1994; Talbot & Hole, 1994). Eutrophication can alter the aquatic ecosystem, inducing diseases, and cause mortalities in exposed organisms (Jasmin et al., 2020; Lananan et al., 2014). As an alternative, bioremediation has been used to eliminating or reducing harmful compounds through its use and biological processing (Jasmin et al., 2020; Divya et al., 2015).

In Mexico, Pacific white shrimp production (*Litopenaeus vannamei*) by its volume is situated in a second-place of the total aquaculture. Its average annual production growth rate in the last ten years has been around 1.67% (CONAPESCA, 2018). However, one of the major problems facing the Pacific white shrimp industry is the low utilization of nutrients supplied through the feed. Generally, only 30% of the nitrogen feed is used by farmed shrimp, while the remaining 70% is discarded or excreted as dissolved form or particles into the water (Troell et al., 1999). A viable strategy to reduce and reuse the discharge of nutrients from the Pacific white shrimp crop is a polyculture’s practice through an Integrated Multi-Trophic Aquaculture (IMTA). The IMTA has the potential to reduce the environmental impact of cultivated species since the wastes are re-utilized as input for another species (Granada et al., 2018; Estim, 2015; Ridler et al., 2007; Chopin et al., 2001). The IMTA systems benefit environments as a result of processes such as bioremediation, and combines economic viability (i.e., cost reduction, diversified performance), social acceptability (i.e., environmental problems, organic quality), and provides a culture with feasible production (Estim, 2015). Among the species of interest where the multitrophic systems have been evaluated are the Pacific white shrimp *Litopenaeus vannamei*, pacific oyster *Crassostrea giga*s and smooth clam *Chione fluctifraga* (Martinez-Cordova and Martinez-Porchas, 2006), seaweed *Gracilaria lemaniformis*, and scallop *Chlamys farreri* (Mao et al., 2009), blue mussel *Mytilus edulis* and Atlantic salmon *Salmo salar* (Reid et al., 2010), rainbow trout *Oncorhynchus mykiss*, Nile tilapia *Oreochromis niloticu*s and seaweeds (*Porphyra, Ulva* and *Gracilaria)*(Pereira et al., 2012), *Eisenia arborea* and red abalone *Haliotis rufescens* (Zertuche-González et al., 2014), Pacific white shrimp *Litopenaeus vannamei* and grey mullet *Mugil cephalus* (Aghuzbeni et al., 2016), Gilthead seabream *Sparus aurata*, grey mullet *Mugil cephalus*, sea urchin *Paracentrotus lividus* and sea cucumber *Actinopyga bannwarthi* (Israel et al., 2019), among others. The previous works have shown that IMTA is a feasible practice for using waste nutrients to produce biomass of cultivated species. However, to achieve maximum use of the feeds and reduce the excess of nutrients that are discharged into the environment, it is necessary to keep evaluating cultured species’ ability to harness the remaining nutrients for its optimal growth.

The grey mullet (*Mugil cephalus*) is a marine fish species whose feeding habits position it as a great candidate for aquaculture. One of the advantages of mullets is that they can be fed from the unused nutrients from the shrimp farming waste (SFW). The grey mullet is a low trophic species with omnivorous habits and can feed on detritus and microflora, it is a euryhaline and eurytherm species. (Aghuzbeni et al., 2016; Moriarty, 1976). Because of those mentioned above, grey mullet is placed as an ideal candidate for its polyculture. Also, grey mullet has a very acceptable marketing value and a great potential for its production in crops (Biswas et al., 2012). In Mexico, there is still no record of its aquaculture production in captivity (CONPESCA, 2018).

To date, most studies evaluating the viability of shrimp-mullet co-cultures have not focused on separately evaluating the contribution of only shrimp farming waste to mullet performance (Hoang et al., 2020; Hoang et al., 2020a; Legarda et al., 2019; Hoang et al., 2018; Aghuzbeni et al., 2015). Most of the studies have evaluated the two species growing them in the same tank, which becomes a real challenge to quantify how much the mullet consumes from the shrimp wastes and how much it consumes from the feed offered to shrimp. This information might be essential to evaluate the contribution of the nutrients obtained from the waste to the mullet’s performance. Therefore, the present study evaluated the use of residual nutrients from Pacific white shrimp farming wastes (i.e., uneaten feed and feces) on the growth performance, digestibility, whole-body proximate composition, and enzymatic activity of juvenile mullet. It also evaluates the incorporation of the Pacific white shrimp farming waste as a co-feed in the juvenile mullet diet as a feeding strategy.

## 2. Material and methods

### 2.1. Ethics statement

All experimental fish used in the present study were handled under the procedures that agreed with the State Committee of Bioethics in Nayarit (Number: 96/CEB/2017) to cause minimal suffering to animals.

### 2.2. Experimental treatments

Four experimental treatments were designed to evaluate the biological performance, feed utilization, apparent digestibility, whole-body proximate composition, and the juvenile mullets’ enzyme activity. The treatments were designed with a commercial feed and a feed based on Pacific white shrimp farming wastes (SFW) offered in different proportions. The proximate composition of the two experimental feeds is shown in table1. The first treatment consisted of offering a generic commercial feed for marine fish (Skretting, marine fish). The second treatment (SFW33) comprised offering 33 % of SFW and 67 % the commercial feed, third treatment (SFW67) comprised offering 67 % of SWF and 33 % the commercial feed, finally, the fourth treatment (EC100) comprised offering the total of the protion of SWF. Every treatment was evaluated in triplicate. The bioassay ended until a significant difference was found in any of the variables evaluated among the treatments, which resulted in 6 weeks.

**Table 1.** Proximate composition of the commercial feed and the shrimp farming waste used for the feeding trial (dry matter basis).

|  | Experimental feeds |  |
| --- | --- | --- |
|  | Commercial feed | Shrimp farming waste |
| Crude protein (%) | 55.6 ± 0.4 | 17.2 ± 0.5 |
| Ether extract (%) | 15.1 ± 0.2 | 1.5 ± 0.3 |
| Ash (%) | 9.1 ± 0.4 | 35.5 ± 0.1 |
| Nitrogen free extract + fiber (%) | 20.4 | 45.8 |

Values are shown as mean (n = 3) ± Standard deviation (SD).

### 2.3. Collection of the Pacific white shrimp farming waste

Shrimp waste was collected from the production Bioengineering Laboratory of the Escuela Nacional de Ingeniería Pesquera from the Universidad Autónoma de Nayarit. The production laboratory contained a culture module consisting of 9 circular geomembrane tanks with a capacity of 80 cubic meters each (10 m diameter; 1 m depth). Each tank contained a density of 100 shrimp per square meter. Shrimps that were cultured in the laboratory were fed with a daily ration of 6% of their biomass. Likewise, they were offered a commercial feed (Previtep Aquamar, Jal, Mexico) with a content of 35% crude protein, 6% lipids, 12% ash, and 31.5% NFE. At the time of waste collection, the temperature was 28 °C, the salinity was 29 ppt, and the dissolved oxygen was 6 mg L. The collection was performed in the morning before the first feeding to avoid collecting fresh feed and to ensure collecting only feces and uneaten feed (i.e., farming wastes).

### 2.4. Preparation of the Pacific white shrimp farming waste pellets

SFW was transformed into feed pellets by adhering and mixing 5% gelatinized starch as a binder. The mixing was performed for 15 minutes with a 7 L^-1^ capacity food mixer (Torrey, model: B7, México). Likewise, about 10% of water was poured into the mixing container to give the mixture’s desired consistency. The mixture was then passed through a meat grinder (Rhino, 1HP model: MOCA-12, México) with a 1/16” diameter die to cold-extrude the pellets. The pellets were dry in an oven at 60 °C for 24 hours. Once the pellets were dried and cooled, they were placed in sealed plastic bags and stored at 4 °C until the feeding trial.

### 2.5. Fish and culture conditions

The mullet’s juveniles were captured from the wild in the estuary El Yugo in Mazatlán, Sinaloa, Mexico. Then, fish were transported to the unit’s laboratory specialized in aquaculture management and innovation in the Centro Nayarita de Innovación y Transferencia de Tecnologia (CENITT) in Tepic, Nayarit, México. Before performing the feeding trial, the mullets were kept for three weeks in laboratory conditions for their acclimatization. A total of 120 juveniles of grey mullet weight of 1.51 g ± 0.0 an average length of 4.6 cm ± 0.2 were randomly distributed in 12 tanks (n = 10 fish per tank).

Each experimental unit consisted of a 45-L tank adapted to a salt-water recirculating system composed of a biofilter of polyurethane balls with a volume of 0.1 cubic meters, a water pump (Lifeguard, Quiet One, 758 GPH, USA), and an air compressor (Boyu Acq-007, China). Water temperature (°C) and dissolved oxygen (mg L^-1^) were measured every day in the early morning using a multi-parameter oxygen meter (YSI model Pro 2030, USA). The concentration of ammonia, nitrites, and nitrates in the tanks were monitored every week with an aquarium set kit (APA®; USA) to keep the values of N-NH3 < 1.0 mg L^−1^, NO2 < 0.3 mg L^−1^, and NO3 < 10.0 mg L^−1^ respectively.

### 2.6. Feeding protocol

The feed was offered to the fish in each tank in a ratio of 4% of their wet weight (g) in accordance with El Sayed (1994) and Wassef et al. (2001). Quantity (g) of the feed ration offered was adjusted weekly after completing each biometry in order always to be offering 4% of the biomass of the tank in the feed. The feed was offered in two portions, the first at 9:00 am and the second 6 hours later.

### 2.7. Growth performance and feed utilization analysis

For monitoring the growth performance of juvenile mullets, fish weight (g) and length (cm) were evaluated weekly using a scale (Adam Equipment, model HCB 1002, USA) and an ichthyometer (Aquatic Eco - Sistems, Inc. FL, USA), respectively. All fish from each experimental tank were sampled and used for every morphometric measurement. The feeding trial’s total feed was determined by adding the daily feed ration offered to each tank.

### 2.8. Calculations of growth performance and feed utilization

Growth performance and feed utilization were evaluated using the following formulas:

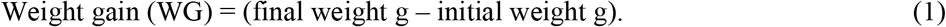

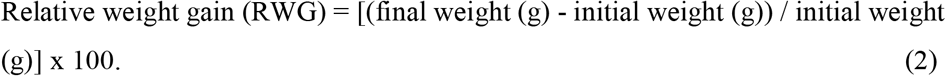

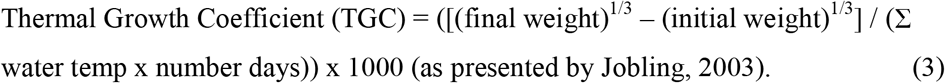

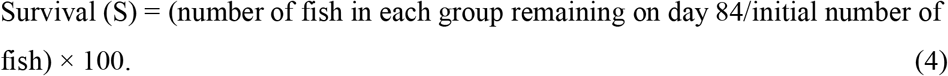

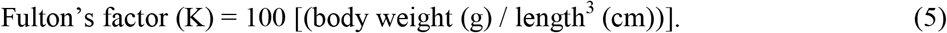

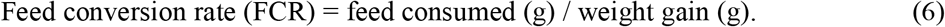

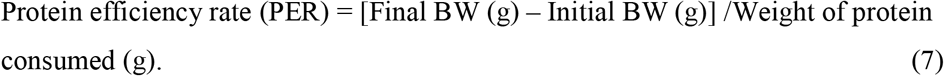

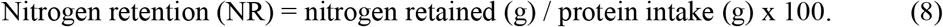

### 2.9. Proximate composition analysis

For the analysis of the whole-body proximate (%) composition, three fish at the beginning of the feeding trial and three fish per tank at the end of the feeding trial were collected by euthanizing them with a clove oil overdose. The samples were frozen and kept at -20° C until evaluating the proximal body composition analysis, initial and final. Five grams of each feed (i.e., commercial feed, shrimp farming waste) were stored in conical tubes and kept at -20° C before analysis for evaluating the feed. The proximate composition of all samples was determined using established methods, according to (AOAC, 1990). Crude protein (N × 6.25) was determined by the micro-Kjeldahl method. Lipids were determined by solvent extraction using petroleum ether in a Soxhlet extractor. Ash was estimated by incinerating the samples for eight hours in a muffle at 550 °C. The nitrogen-free extract (NFE) was calculated by subtracting the added percentage of protein + fat + ash from 100%.

### 2.10. Apparent digestibility analysis

The fecal material for evaluating the apparent digestibility coefficients was collected daily. The collection started seven days after running the feeling trial until enough material was obtained to perform digestibility analysis. Feces were collected from the bottom of the tank. It used a 20-cm long and 2-mm wide PVC siphon immediately after the feces’ release was observed (approximately 1hour after the feed was offered) to avoid leaching. The collected samples were dry at 60 °C in an oven for 24 h to extract humidity. Then they were stored at −4 °C in conical-bottom polypropylene 50 ml capacity tubes for later analysis.

Determination of the ADC’s of the dry matter and crude protein were calculated using the insoluble ash in hydrochloric acid as an internal marker. The estimation of the values was assessed according to the method of Montaño-Vargas et al. (2002).

The acid-insoluble ash (AIA) was calculated as follows:

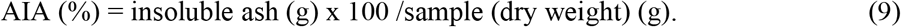

The ADC’s of the dry matter and the CP of the diets were estimated as follows:

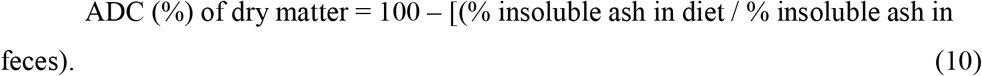

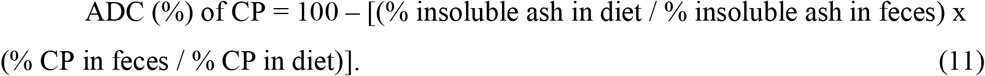

### 2.11. Enzyme activity

For enzymatic analyses, two juvenile mullets from each experimental tank were collected and lyophilized before analysis. The whole-body of juvenile mullets was processed since they were too small to be dissected and obtain enough sample material for their analysis. The enzyme extract was obtained by placing the samples in a conical 15 ml tube, previously cooled, to be homogenized with a tissue grinder (POLYTRON® PT 1200, Kinematica AG, Switzerland), in 10 ml distilled water at 4 ° C. Once the extract was homogenized, it was centrifuged at 16000 g for 30 minutes, at 4 ° C, and the supernatant was collected and stored in 0.5 ml aliquots at -80 °C. Aliquots, once thawed, were used within 48 h and kept refrigerated at 4 °C. Total activity per fish was estimated as units of absorbance per mg (U mg^-1^). The procedures were adapted to perform the spectrophotometric measurements on a plate reader Varioskan Flash (Thermo Scientific), and the data were processed in the software SkanIt RE 2.4.5. A blank was included in each enzyme determination using distilled water instead of the homogenate.

The trypsin activity was measured according to the method of Erlanger et al. (1961). The procedure consisted of using one mM of BAPNA (Nα-benzoyl-DL-arginine-p-nitroanilide hydrochloride, Sigma B-4875) substrate dissolved 500 µL of DMSO in 50 mM Tris-HCl buffer, 20 mM CaCl2, pH 8.2 at 37 °C by 30 min. The reaction was stopped with 30 % acetic acid, and the absorbance was recorded at 410 nm after 10 minutes of stabilization.

The chymotrypsin activity was according to the method of Hummel (1959), as modified by Applebaum et al. (2001), using 0.56 mM of BTEE (N-Benzoyl-L-tyrosine ethyl ester, Sigma 13110-F) as a substrate in 100 mM Tris-HCl buffer, 25 mM CaCl2, pH 7.8 and methanol 2.5 % (v/v) at 37 °C. The reaction was recorded every minute for 30 min at 256 nm in a 96-well quartz bottom plate.

The total alkaline protease activity was measured according to Sarath et al. (1989) using 2 % of casein (Sigma, C5890) as a substrate in 50 mM HCl-Tris buffer, ten mM CaCl2, 9.0 pH at 37 °C. The reaction was incubated for 10 min at 37 °C and stopped with 10 % trichloroacetic acid. Finally, the samples were centrifuged (21000 g, 5 min, 4 °C), and the supernatant’s absorbance was recorded at 2800 nm. According to Worthington Biochemical Corporation (1993), amylase activity was measured using as substrate 1 % of starch (Sigma, S9765). The starch was mixed in 20 mM sodium phosphate buffer, six mM NaCl and 6.9 pH. The homogenate and the buffer + substrate were mixed at intervals of time, incubate at 25°C, and at 3 min, 1 % dinitrosalicylic acid was added as a colour reagent. After, all tubes were incubated in a boiling water bath for 5 minutes. Afterward, tubes were cooled at ambient temperature and mixed to read absorbance at 540 nm. A standard maltose curve was used to calculate the micromoles of maltose released in each sample by the regression equation.

All the samples were diluted ten times in a final volume reaction of 1.2 ml to allow readings within the standard range.

The lipase activity was estimated according to the method of Gjellesvik et al. (1992) using 0.56 mM of 4-Nitrophenyl myristate (Sigma 70124) as substrate dissolved in 0.5 mL DMSO. The reaction conditions were 150 mM Tris-HCl buffer, 15 mM sodium taurocholate, pH 8.5 at 37 °C. Also, the reaction was recorded every minute by 30 min at 405 nm. Pepsin activity (i.e., acid proteolytic activity) was estimated according to Sarath et al. (1989), using 1 % of hemoglobin as substrate (Spectrum Chemical, HE120) in 200 mM, pH two at 37 °C. The reaction was incubated for 10 min and stopped with 5 % trichloroacetic acid. The samples were centrifuged (21000 g, 5 min, 4 °C), and the absorbance of the supernatant was recorded at 280 nm. In all cases, one unit of enzyme activity was defined as the amount of enzyme required to cause an increase of 1 unit of absorbance (Lazo et al., 2000). Activities were expressed in units per gram of wet tissue.

### 2.12. Statistics

All variables’ values were analyzed using a completely randomized design and presented as averages (n = 3) ± standard deviation (SD). All data were evaluated for their normality with a Komologorov-Smirnov test and their homoscedasticity with a Levene test. A one-way analysis of variance (ANOVA) test was applied to determine possible statistical differences among treatments and a Tukey test when differences were found (P < 0.05). Before analysis, all percentages (%) data were arcsine transformed. A Student t-test with a significance level of 0.05 was used to find a significant difference in the apparent digestibility coefficients between the commercial feed and the shrimp farming waste. All the statistical tests were performed using the STATISTICA® program, version 7.

## 3. Results

During the feeding trial, no signs of disease were observed in the juvenile mullets. The temperature registered an average of 25.6 + 0.4 ° C regarding water quality parameters, the salinity of 28.6 ± 1.1, the pH of 7.8 ± 0.1, and ammonia maintained an average value of 0.1 ± 0.1 mg L^-1^, nitrites of 0.01 ± 0.0 mg L^-1^, nitrates of 0.01 +0.0 mg L^-1^, and dissolved oxygen at 5.7 ± 0.24 mg L^-1^.

### 3.1. Growth performance and feed utilization

At the end of the feeding trial, the juvenile mullets of treatment SFW0 registered the highest value in FW, WG, RWG, and TGC, with a significant difference (P < 0.05) among the treatments (Table 2). However, although juvenile mullets from the SFW100 treatment recorded the lowest growth performance values, the feeding trial achieved an RWG value of 13.16 ± 5.3. Likewise, they had an increase of 0.2 g of the WG. At the end of the feeding trial, there were no differences in the survival percentages among the treatments. Regarding the K values, the SFW0 and SFW100 treatments had the highest values, being different (P < 0.05) from the SFW33 and SFW67 treatments’ values. Experimental feed was never observed at the bottom of the tanks during feeding, indicating that it was always consumed (4% of biomass). However, significant differences (P < 0.05) were observed among treatments on the feed utilization values.

**Table 2.**
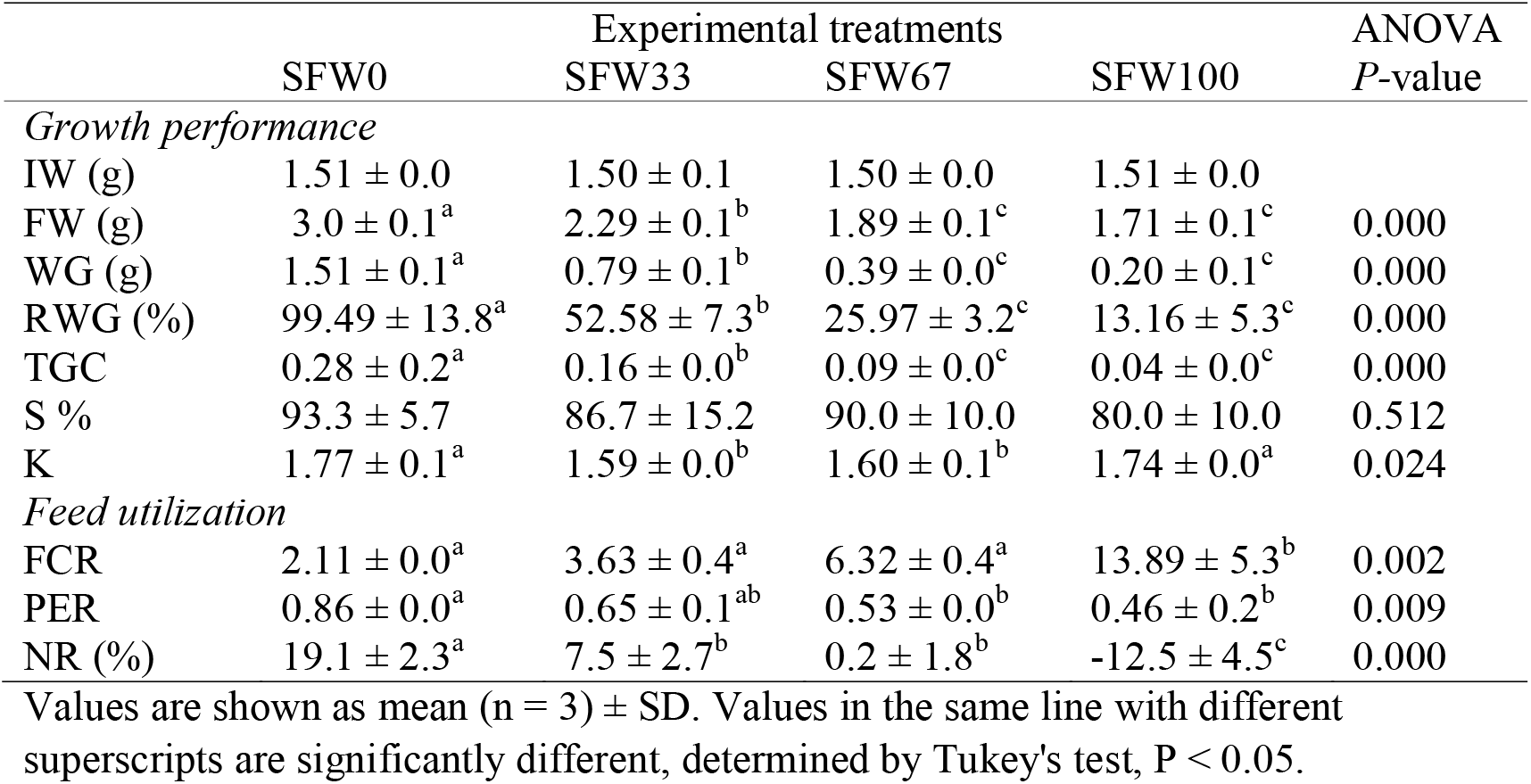
Growth performance and feed utilization of juvenile grey mullet at the end of the feeding trial.

The juvenile mullets of treatment SFW100 registered the lowest value of FCR. However, although the feed offered to juvenile mullets of treatments SFW0, SFW33, and SFW67 contained different CP levels in the feed, the FCR values among treatments mentioned did not show a significant difference (P < 0.05). The PER values were significantly higher in treatment SFW0. However, the remained treatments did not show differences among them. Although the feed received among them had different levels of crude protein. Juvenile mullets from the SFW100 treatment registered the lowest NR value, even with negative values.

### 3.2. Proximate composition of the whole-body

The proximate composition (%) values were different among treatments, as well as the initial sample. Mullets of treatment SFW0 registered the lowest values of moister. However, the rest of the treatments did not show statistical differences, even with the initial sample. The CP presented the highest values in the initial sample and the juvenile mullets of the SFW0 treatment. Also, it was observed that as the SFW ratio increased in the treatments, the CP content decreased in the juvenile mullets, registering the lowest value in the juvenile mullets of the SFW100 treatment. Regarding the ash content, no significant differences (P < 0.05) were found among the treatments, not with the initial sample. The rest of the proximate composition values are found in Table 3.

**Table 3.** Proximate composition (wet weight) of the whole-body of juvenile grey mullet at the end of the feeding trial among treatments and at the beginning of the feeding trial.

|  | Initial | Treatments |  |  |  | ANOVA |
| --- | --- | --- | --- | --- | --- | --- |
|  |  | SFW0 | SFW33 | SFW67 | SFW100 | P-value |
| Moisture (%) | 72.1 ± 1.06<sup>ab</sup> | 69.07 ± 1.68<sup>b</sup> | 73.05 ± 1.90<sup>ab</sup> | 74.51 ± 6.39<sup>ab</sup> | 78.07 ± 5.98<sup>a</sup> | 0.022 |
| Crude protein (%) | 18.04 ± 0.14<sup>ab</sup> | 20.66 ± 1.57<sup>a</sup> | 16.61 ± 2.2<sup>bc</sup> | 15.42 ± 0.18<sup>bc</sup> | 13.79 ± 0.29<sup>c</sup> | 0.000 |
| Ether extract (%) | 3.53 ± 0.31<sup>ab</sup> | 4.88 ± 0.62<sup>a</sup> | 3.26 ± 1.05<sup>ab</sup> | 2.75 ± 1.16<sup>bc</sup> | 1.05 ± 0.27<sup>c</sup> | 0.002 |
| Ash (%) | 5.26 ± 0.31 | 5.29 ± 0.22 | 5.99 ± 0.46 | 6.38 ± 0.63 | 5.77 ± 0.82 | 0.110 |
| Nitrogen free extract + fiber | 1.05 | 0.10 | 1.09 | 0.94 | 1.32 |  |

Values are shown as mean (n = 3) ± SD. Values in the same line with different superscripts are significantly different, determined by Tukey's test, P < 0.05.

### 3.3. Apparent digestibility coefficients

In the present study, the apparent digestibility coefficients of commercial feed and shrimp farming wastes in gray mullets were evaluated since the 4 treatments were created from these two experimental feeds. The results showed that the highest (P < 0.05) values of the apparent digestibility coefficients with commercial feed were observed, both for the feed’s dry matter and protein. However, the juvenile mullets fed with the shrimp farming wastes achieved a protein digestibility value of 41.29%. The rest of the values of the digestibility coefficients are presented in table 4.

**Table 4.** Apparent digestibility coefficients of the dry matter and crude protein of the commercial feed (CF) and the shrimp farming wastes (SFW) of juvenile grey mullet.

|  | Experimental feeds |  | T-test |
| --- | --- | --- | --- |
|  | CF | SFW | *P*-value |
| Dry matter | 77.18 ± 6.08 | 35.42 ± 7.60 | 0.001 |
| Protein | 91.37 ± 1.98 | 41.28 ± 4.33 | 0.000 |

Values are shown as mean (n = 3) ± SD.

### 3.4. Enzyme activity

Although the feed offered in the experimental treatments had different levels of protein and lipids, the evaluated values of the activity of digestive enzymes (i.e., alkaline proteases, trypsin, chymotrypsin, amylases, and lipases) did not show significant differences among treatments (P > 0.05). The results are shown in Table 5.

**Table 5.** Digestive enzyme activity (U mg fish^-1^) of alkaline proteases, trypsin, chymotrypsin, amylases, and lipases of juvenile grey mullet fed with the experimental treatments.

|  | Treatments |  |  |  | ANOVA |
| --- | --- | --- | --- | --- | --- |
|  | SFW0 | SFW33 | SFW66 | SFW100 | *P*-value |
| Alkaline proteases | $0.275 \pm 0.05$ | $0.243 \pm 0.05$ | $0.263 \pm 0.04$ | $0.256 \pm 0.05$ | 0.8949 |
| Trypsin | $0.376 \pm 0.28$ | $0.471 \pm 0.26$ | $0.257 \pm 0.13$ | $0.385 \pm 0.25$ | 0.7638 |
| Chymotrypsin | $1.388 \pm 0.08$ | $1.617 \pm 0.19$ | $1.606 \pm 0.03$ | $1.630 \pm 0.01$ | 0.0708 |
| Amylases | $0.011 \pm 0.004$ | $0.010 \pm 0.003$ | $0.007 \pm 0.003$ | $0.008 \pm 0.002$ | 0.6601 |
| Lipases | $0.251 \pm 0.02$ | $0.272 \pm 0.11$ | $0.232 \pm 0.06$ | $0.252 \pm 0.101$ | 0.9494 |

Values are shown as mean ( $n = 3$ ) ± SD.

## 4. Discussion

As aquaculture grows, it is necessary to maintain a sustainable practice to reduce the environment’s negative impact (FAO, 2020; Granada et al., 2018). Information related to the use of residual nutrients from aquaculture species is necessary to practice sustainable aquaculture and reduce excess nutrients discharged into the environment. In the present study, the use of residual nutrients by juvenile mullet for its growth performance was evaluated. At the end of the feeding trial, a notable difference in the growth variables among treatments was observed. The juvenile mullets of SFW0 treatment obtained higher (P < 0.05) values on the growth variables (i.e., FW, WG, RWG, and TGC). Higher growth in juvenile mullets was expected due to the protein and lipid content of the feed offered (100 % commercial feed) in the SFW0 treatment,compared to the treatments that were offered different percentages (33, 66, 100%) of SFW. In the present study, crude protein and lipid in the SFW resulted in 17.2 and 1.5% content, respectively. In a similar study, Israel et al. (2019) collected the wastes from a gilthead sea bream farm, *Sparus aurata*; the nutritional content of the waste they collected resulted in 18.7% crude protein and 6% lipids. It is noteworthy that the nutrient content of the farming wastes can vary depending on the digestibility of the species cultivated (Galasso et al., 2017; Herath and Satoh, 2015). Digestibility depends on biological, environmental, and dietary factors (Sugiura, 2000). Likewise, it was noted to observe a decrease in the mullet’s biological performance as the percentage of SFW when the percentage of feed increased. These results suggest that if the juvenile mullet feed only on SFW, they will not obtain enough nutrients to obtain optimal growth. Since a protein level of 30% has been determined in the feed to cover the mullets’ protein requirement (Talukdar et al., 2020) and a level of 6% lipids (De et al.,2011). However, Hoang et al. (2018) determined that a maximum of 10% biomass of mullets (*Mugil cephalus*) should be cultivated concerning Pacificshrimp’s (*Litopenaeus vannamei*) cultivated biomass to obtain greater productivity and utilization of nutrients.These data were determined when the two species were kept in the same culture tank.Hoang et al. (2020) observed in a shrimp-mullet (*Litopenaeus vannamei-Mugil cephalus*) co-culture that the shrimp was cultivated with the mullets kept in separate cages, the mullets’ growth performance obtained low values. Similar results were reported by Borges et al. (2020) evaluating shrimp-mullet (*Litopenaeus vannamei-Mugil liza*) co-culture with biofloc technology. The authors reported that when mullets are grown in separate tanks from shrimp, the mullets’ biological performance is considerably reduced compared to when they are grown in the same tank. The results mentioned above agree with the obtained in the present study, noticing that when the mullets were co-fed with SFW, a decrease in growth performance was observed. The data suggest that it is not enough to feed them only with SFW for optimal mullet growth. It is necessary to supplement feeding with formulated feed or some source of extra nutrients like biofloc, as suggested by Legarda et al. (2019).

Regarding the feed utilization values, as the SFW increased as feed, the FCR values increased, and the PER and NR values decreased. These results are due to the lower amount of nutrients (i.e., protein, lipids) that SFW contained. Several studies that evaluated the mullets’ biological performance fed with the shrimp waste did not report the mullets’ FCR values (Hoang et al., 2020a; Hoang et al., 2020b; Borges et al., 2020). Probably because of the difficulty of measuring the amount of waste that mullets consumed. Legarda et al. (2019) reported lower FCR values than the present study. However, that study used biofloc technology, which provides considerable nutrient content that utilized mullets to grow. A decrease in nitrogen retention (NR) values was observed as the SFW was increased in the feed, registering significant differences among the treatments. At the end of the feeding trial, a more significant reduction in the NR value was observed in the SFW 100 mullets. This result could be because of the lack of feed nitrogen, and mullets had to catabolize nitrogen from the body to produce their energy. Enzymes for catabolism and amino acid synthesis occur in each tissue. The catabolism process involves the deamination resulting in a carbon skeleton that can be channeled into the tricarboxylic acid cycle, where it is either oxidized that can be oriented towards gluconeogenesis via pyruvate carboxylase (Bequette, 2003). Due to catabolic reactions, energy is provided for different metabolic actions such as mechanical work, transport, and anabolic activity such as the synthesis of carbohydrates, proteins, and fats (McDonald et al., 2011).

The SFW percentage in the treatments affected the mullets’ proximal whole-body composition at the end of the feeding trial. The reduction of protein and lipids in the whole-body was notable as the SFW percentage in the feed ration increased, showing the lowest values with the SFW100 treatment mullets. This result can be explained due to the low protein and lipid content in SFW. The lack of nutrients in the feed leads to a nutritional deficiency, deteriorates the fish’s health, and affects the composition of nutrients in the tissues (Lall and Dumas, 2015; NRC, 2011). Likewise, protein deficiency in feed reduces growth due to the extraction of proteins from less vital tissues (i.e., muscle) to prioritize the most vital functions (Wilson, 2002). A similar result was observed by Talukdar et al. (2020), whom found a higher protein accretion in the mullets’ carcass when they utilized 30 and 32% protein levels in the feed. Biswas et al. (2012) also found differences in whole-body protein and lipid composition (%) of the mullets. The proximal composition (% in dry matter) of the experimental feed was 27.5 CP and 5.2 L. They mention that the formulated feed is accepted rapidly and contains a balance of nutrients, helping the mullets’ performance. However, Gisbert et al. (2016) found no differences in mullet fingerlings’ proximate composition evaluating different weaning diets. It is worth mentioning that although they used different protein sources in the diets, they remained isoproteic and isolipidic, covering the nutritional requirement of the mullets. Also, De et al. (2012) found no effect on carcass protein composition in mullets fed with different protein levels (i.e., 20, 25, 30, and 35 % of CP, in dry matter), which might be because the mullets had enough protein in the diet and did not need to use tissue protein as an energy source.

Although mullets have a notable ability to feed on sediments or detritus (Whitfield et al., 2012), in the present study, we found an evident low digestibility of the dry matter and protein when feeding on SFW (i.e., below 50 %). Waste farms usually decompose very fast (Galasso et al., 2017); the feed composition is one of the factors that affect organisms’ digestibility (Sugiura, 2000), as well as the high amount of ash in the feed (NRC, 2011). However, one of the essential nutritional characteristics of mullets is that they have the ability to feed on the organic matter of the sediment (Silva, 1980), making the mullets an ideal species for polyculture and obtaining nutrients from farm waste (Lupatsch et al., 2003). Israel et al. (2019) found low coefficients of apparent digestibility of dry matter and protein in mullets, feeding them with seabream (*Sparus aurata*) farm waste. The authors mention that for farm waste to have nutritional value for mullets, the waste must remain in the bottom sediment to accumulate nutrients and as well as produce microbial biomass and contribute to the nutritional value. The present study agrees with what was commented by Israel et al. (2019) since a lack of nutritional value was observed in the SFW, negatively affecting the mullets’ growth performance. The result viewed from the sustainability of aquaculture and reduction of the environmental impact may be favorable. The reuse of shrimp farming wastes as feed for mullets may be a viable practice. However, to obtain crop production, it would be necessary to feed the mullets with feed formulated to satisfy the species’ nutritional requirements. Our results showed that there were no significant differences in the activity of digestive enzymes between the treatments. Generally, fish are able to adjust the secretion of pancreatic digestive enzymes concerning the level of feed and its quality (Buddington et al., 1997). Omnivorous fish, such as grey mullets, generally have a high activity of many of the digestive enzymes and are able to use a wide range of food sources (NRC, 2011). It could have been that at the time of the collection of the fish sampled, and digestive enzymes were being secreted to be ready for feed consumption, regardless of the source, be it commercial feed or shrimp farming wastes, registered no significant difference among treatments. In subsequent studies, it is recommended to evaluate the enzymatic activity per organ of fish to eliminate the possible reading of non-digestive enzyme activity.

## Conclusion

The present study suggests that mullets are capable of utilizing residual nutrients from shrimp farming waste. However, to have optimal growth performance, the addition of an alternative feed source is necessary. Utilizing residual nutrients from shrimp farm waste as a feed source for mullets will contribute to sustainable aquaculture and reduce the environment’s impact due to the excess of residual nutrients discharged. However, to meet nutritional requirements, it is recommended that the mullets be fed with formulated feed so that the protein and lipid content in the whole-body of the mullets do not decrease and increase growth performance under growing conditions.

## Acknowledgments

Thanks to the grant from Consejo Nacional de Ciencia y Tecnologia (CONACyT) for the support of the postdoctoral stay (Convocatoria: Estancias Posdoctorales 1er Año 2019 – 1). Thanks to the State Council of Science and Technology (COCYTEN), of the government of Nayarit, Mexico, for the partial financing of the experiments, though the project “Ordenamiento y Tecnificacion del Cultivo de Camaron en Nayarit: Primera Fase.” This study was partially funded by the Consejo Nacional de Ciencia y Tecnologia (CONACYT-SEP-CB-2016-284167) coordinated by Dr. Zohar-Zatarain.

